# ScanExitronLR: characterization and quantification of exitron splicing events in long-read RNA-seq data

**DOI:** 10.1101/2022.03.25.485864

**Authors:** Joshua Fry, Yangyang Li, Rendong Yang

## Abstract

**Summary:** Exitron splicing is a type of alternative splicing where coding sequences are spliced out. Recently, exitron splicing has been shown to increase proteome plasticity and play a role in cancer. Long-read RNA-seq is well suited for quantification and discovery of alternative splicing events; however, there are currently no tools available for detection and annotation of exitrons in long-read RNA-seq data. Here we present ScanExitronLR, an application for the characterization and quantification of exitron splicing events in long-reads. From a BAM alignment file, reference genome and reference gene annotation, ScanExitronLR outputs exitron events at the transcript level. Outputs of ScanExitronLR can be used in downstream analyses of differential exitron splicing. A companion tool, AnnotateExitron, reports exitron annotations such as truncation or frameshift type, nonsense-mediated decay status, and Pfam domain interruptions. We demonstrate that ScanExitronLR performs better on noisy long-reads than currently published exitron detection algorithms designed for short-read data.

**Availability and Implementation:** ScanExitronLR is freely available at https://github.com/ylab-hi/ScanExitronLR and distributed as a pip package on the Python Package Index.

**Supplementary Information:** Supplementary data are available at *Bioinformatics* online.

## 1 Introduction

An exitron is a region within an annotated coding exon that is spliced out like an intron. Exitrons are unique in that they possess both protein-coding and intronic potential while also possessing canonical splice-site signals (e.g. GT-AG). Originally described in *Arabidopsis thaliana*, exitrons have been shown to increase plant proteome diversity and plasticity (Marquez et al, 2015), mediate responses to heat stress (Cecchini et al, 2022) and create novel gene isoforms (Cheng et al, 2020; Aliperti et al, 2019). In humans, exitrons can alter cancer driver genes, promote tumor progression and be a potential source of neoantigens (Wang et al, 2021). Because exitrons were discovered relatively recently, nomenclature has not yet settled and they have also been called ‘cryptic introns’ (e.g. Dean et al, 2020) or ‘intron retention loss’ (e.g. Ringeling et al, 2022).

Long-read sequencing, though more error prone than short-read sequencing, is in a better position to identify novel splicing isoforms (Amarasinghe et al, 2020), such as those containing exitrons. However, there are currently no tools available for exitron detection in long-read sequencing. Moreover, novel splice site detection within noisy long-reads presents its own challenges. The higher error rate often leads to imprecision in the exon-intron boundary. To combat this, long-read aligners such as Minimap2 (Li, 2018) can use transcriptome annotations to preferentially align junctions within reads to annotated exon borders. However, this sometimes causes exitrons to be misaligned as alternate 3′ or 5′ splice sites (Supplementary Figure S1, S2).

To solve these issues, we present ScanExitronLR, an application for the discovery and annotation of exitron splicing events in long-reads. In addition to overcoming the lower accuracy of long-reads, ScanExitronLR uses LIQA (Hu et al, 2021) to quantify exitron expression at the transcript level.

## 2 Description of the Algorithm

### 2.1 Implementation details

The input to ScanExitronLR is a BAM alignment file, along with a reference genome and reference gene annotation file (Fig 1a). For alignment, we suggest using Minimap2 (Li, 2018) and supplying a junction bed file for more accurate spliced alignments. ScanExitronLR first identifies introns within the BAM alignments and intersects them with annotated exons to find introns completely contained within an annotated exon. Because unannotated splice-sites may be noisy, we add a small amount of nucleotide *jitter* (10 as default) and treat all exon borders as fuzzy. Thus, a splice-site will be considered novel only if it occurs +/- *jitter* away from annotated splice-sites. We then assign exitrons into clusters such that every exitron within the cluster is no more than *2 * jitter* away from each other. For each cluster, we nominate a consensus exitron that (1) has canonical G[T/C]-AG splice sites and (2) has the highest number of supporting reads. We then realign all exitrons within the cluster to the consensus exitron, thus treating the cluster as one splicing event.

**Figure 1.**
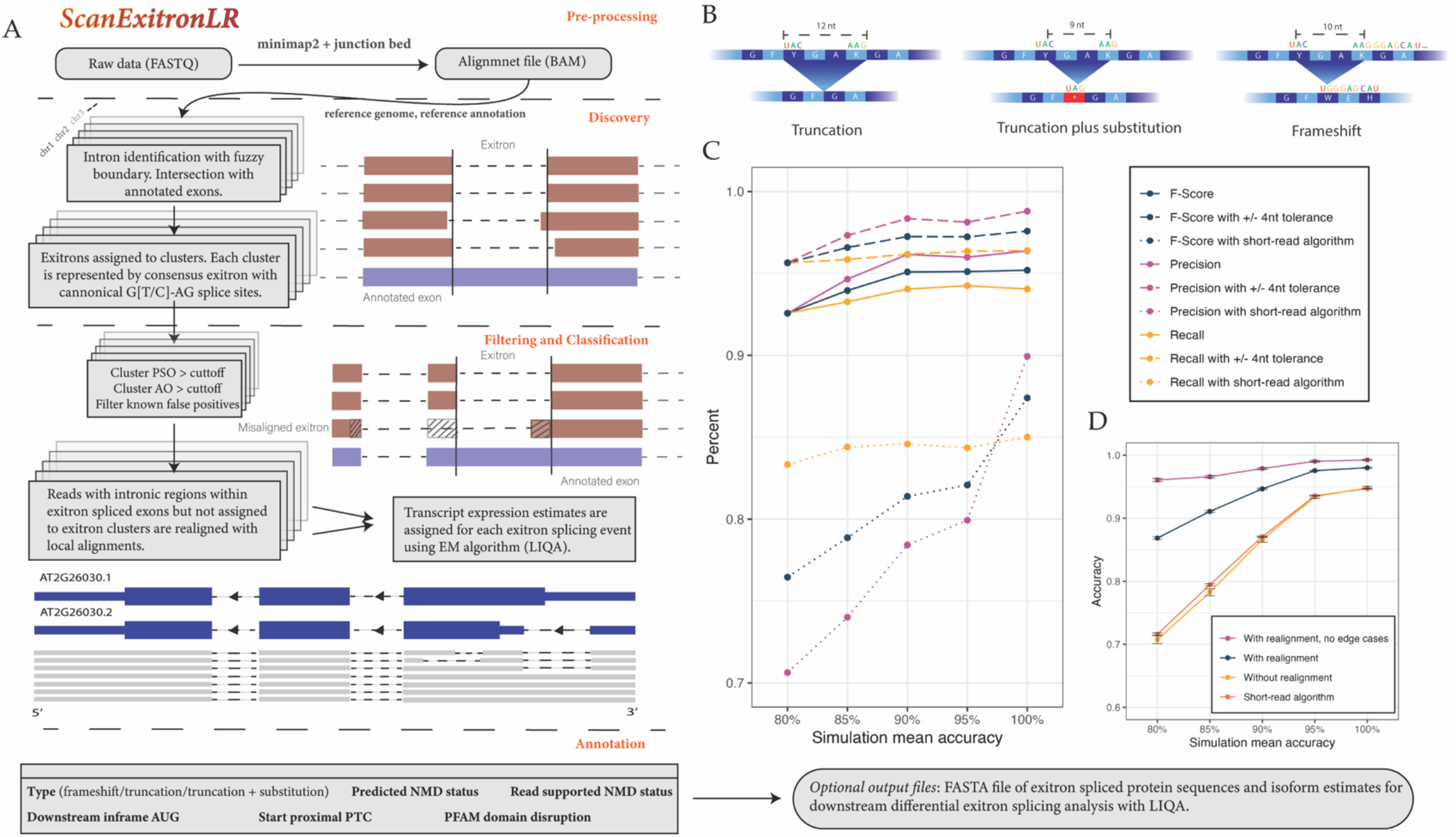
(A) Description of ScanExitronLR and companion tool AnnotateExitron. (B) Schematic representation of the difference between truncation, truncation plus substitution and frameshift inducing exitron splicing events. (C) Simulation results from ONT long-reads at 80%, 85%, 90%, 95% and 100% mean read accuracy levels. (D) Mean accuracy for each detected exitron, defined as (algorithm reported AO)/(ground-truth AO). Error bars represent 95% confidence intervals.

After clustering, we filter exitrons based on AO (number of reads supporting the exitron), PSO (percent spliced out, a measure of the percentage of transcripts for which the exitron is spliced), and, optionally, cluster purity (measured as the proportion of reads with the consensus exitron splice-sites). ScanExitronLR has the option to execute calling and filtering processes for each chromosome in parallel, and we encourage users to use as many cores as possible.

For every exitron that passes filtering, we examine whether reads aligning to the exitron’s position which were not called in the previous step could have harbored misaligned exitrons (Supplementary Figure S2a,b; Fig 1a). Using two rounds of local alignments, a read is realigned as supporting an exitron if, first, the exitron sequence does not appear in the region of the read aligned to the annotated exon and, second, a top scoring local alignment contains a gap with roughly the correct length and flanking splice sequences (Supplementary Methods).

Once exitrons are called, filtered, and realigned, we separate exitron spliced reads from unspliced reads and run LIQA (Hu et al, 2021) for exitron specific transcript quantification. We also provide a companion tool, AnnotateExitron, which takes the output of ScanExitronLR and determines the type of each exitron: frameshift, truncation, or truncation plus substitution which have the potential to substitute a novel amino acid or even stop codon into the truncation site (Fig 1b). We also report predicted and read supported nonsense-mediated decay status as well as other NMD related features (Supplementary Methods). In addition, we report Pfam protein domain disruption, for either *Arabidopsis* or human protein domains. The user can optionally save a FASTA file of exitron spliced protein sequences and isoform estimates for further downstream splicing analysis.

### 2.2 Simulation and real data benchmarking

In order to assess the performance of ScanExitronLR, we performed simulation experiments with simulated long-reads at mean read accuracies of 80%, 85%, 90%, and 100% using PBSIM2 (Ono et al, 2021; Supplementary Methods). Because there are currently no published tools to detect exitrons in long-reads, we instead compared the performance of ScanExitronLR with a previously published tool, ScanExitron, which is designed to detect exitrons in short-read RNA-seq (Wang et al, 2021a; Wang et al, 2021b).

Our simulation results show that the performance of ScanExitronLR is stable across all accuracy levels (Fig 1c). At 80% read accuracy the precision and recall is 92.57% and 92.58% respectively, while at 100% accuracy it is 94.05% and 96.38%. In contrast, the short-read algorithm precision is significantly lower for noisy long-reads, 70.62% at 80% accuracy and 89.93% at 100% accuracy. The 100% to 80% percent difference of the F-score for the short-read algorithm is 10.95% compared to only 2.62% for ScanExitronLR. This shows that our algorithm is indeed able to correct for the errors unique to long-read sequencing. Many of the errors made by our algorithm are mistaken splice-sites. Across all accuracy levels, the F-score increases by around 2% if we allow the true splice-sites to be within +/- 4 nt of our reported splice-sites.

We also computed the accuracy of each true exitron detection event as (algorithm reported AO)/(ground-truth AO) (Fig 1d). Without our realignment step, ScanExitronLR is just as accurate as the short-read algorithm. However, with realignment, the accuracy is significantly increased, especially at low read accuracy levels. We observed that many of the exitron spliced reads not detected by ScanExitronLR were due to faulty alignments when the exitron splicing occurred close to the exon border (Supplementary Figure S2b). Excluding edge cases where an exitron occurs within 50 nt of an exon border, ScanExitronLR is more than 95% accurate across all read accuracies.

Lastly, we compared long-read exitron quantification predicted by ScanExitronLR with short-read predictions by ScanExitron on direct RNA ONT data from three cancer cell lines from the Singapore Nanopore Expression Project (SGNex v0.2) and ScanExitron on Illumina short-read replicates (Chen et al, 2021). We found ScanExitronLR achieved stronger correlation of PSOs between short-read and long-read compared to ScanExitron (Supplementary Figure S3b,c; Supplementary Methods)

## 3 Example

We ran ScanExitronLR on a recently published direct RNA sequencing dataset of *Arabidopsis* samples (Zhang et al, 2020). With an AO cutoff of 2 and PSO cutoff of 0.05, we found 172 exitrons across four biological replicates, two buds and two flowers (Supplementary Methods). As an example, we identified an exitron with a length of 90 nt in gene AT2G26030, an F-box containing protein. This exitron was not detected in the original *Arabidopsis* exitron dataset through short-read RNA-seq analysis (Marquez, et al 2015), though longer exitrons were found in this same gene. Because ScanExitronLR can identify exitrons at the transcript level, using the output of ScanExitronLR in downstream analysis we were able to discover that, in bud samples, exitrons within this gene are differentially expressed in a shortened transcript, AT2G26030.2, with alternative start codon (*p* = 1.88×10^−4^, chi-squared test; Fig 1a). Interestingly, this shortened transcript splices out the F-box binding domain. Thus, this exitron is most likely associated with alternative functions of this gene--an insight one could not obtain without transcript level quantification of exitron splicing events.

