## Supplemental Methods and Figures for "ScanExitronLR: characterization and quantification of exitron splicing events in long-read RNA-seq data"

***Supplementary Methods and Figures***

### S1 Realignment Procedure

After identifying an exitron splicing event that passes filtering conditions, ScanExitronLR attempts to identify misaligned reads that could harbor an exitron. For every called exitron, we identify reads that align to the exitron's position and contain an intron within the annotated exon that is being spliced. For each of these reads we conduct two rounds of local alignments. First, we align the exitron sequence to the read sequence (match score = 2, mismatch score = -1, gap opening penalty = -2, gap continuation penalty = -1). If the read indeed does harbor the exitron, then the alignment score will be low. We normalize the score by the length of the read sequence such that 0 is the worst possible alignment score and 1 is the best possible alignment score. We then only consider reads with normalized alignment score < 0.7, indicating that the exitron sequence is not present anywhere in the read. To see if the read plausibly contains the exitron's splice sites, we align (1) the portion of the read mapped to the annotated exon that contains the exitron and (2) the reference exon sequence (match score = 4, mismatch score = -2, gap opening penalty = -2, gap continuation penalty = 0). We then consider the top scoring alignments. If two conditions are met, we add the read towards the exitron AO and PSO. First condition: there is some top scoring alignment with a total number of gaps +/- (*length_of_exitron* + 10 + *length_of_exon* * 0.05). The idea is that the best local alignments should only have as many gaps as the length of the exitron sequence, plus some jitter (10) and plus gaps perhaps opened by read mismatches, deletions or other effects, proportional to the length of the exon (*length_of_exon* * 0.05, chosen as a heuristic). Second condition: there is some gap within the top local alignment that has the two flanking nucleotide sequences of the exitron. For example, if the exitron is ..AT|GT--------AG|GC, then the top local alignment must have a gap of the form ..AT--...--GC.

Thus, a realigned exitron is called if the exitron sequence, first, does not appear in the region of the read aligned to the annotated exon and, second, contains a gap with roughly the correct length and flanking splice sequences.

### S2 Exitron annotations

The output of ScanExitronLR is an .exitron file describing the locations and quantities of each exitron detected. The companion tool, AnnotateExitron, takes an .exitron file, genome reference and reference gene annotation file and, optionally, the original BAM alignment file, and outputs various features of each exitron. For each exitron, ScanExitronLR reports each transcript with at least 10% abundance. Because exitron features depend on which transcript is being spliced, AnnotateExitron reports annotation features for each transcript reported by ScanExitronLR. First, it reports either a truncation, truncation plus substitution, or frameshift (Supplementary Figure 3a). Both truncations and truncations plus substitutions preserve the reading frame; however truncations plus substitutions have the potential to substitute a novel amino acid or even stop codon into the truncation site. Frameshift exitrons do not preserve the reading frame and have the potential to activate out of frame premature stop codons (PTC) and trigger nonsense mediated decay (NMD). NMD triggering requires that there occurs an exon-exon junction at least 50 nt downstream of the stop codon. AnnotateExitron determines if this condition is true based on the exitron's gene transcript and reports this as the 'predicted NMD status'. If a BAM alignment file is provided, it also checks whether the exitron spliced reads empirically support NMD triggering by having an intron at least 50 nt downstream of the frameshift-induced PTC.

According to a recent study of features that modulate NMD[1], NMD efficiency is lowered if there is a downstream in-frame AUG start codon after the PTC. This is because the translational machinery may reinitiate at the downstream start codon and evade NMD. Thus we report the presence of a downstream in-frame AUG start codon. In addition, transcripts are insensitive to NMD if the PTC occurs close to the transcript start codon. Thus we report whether the PTC occurs within 200 nt of the start codon. If either of these conditions are true for a given exitron, it is evidence that the frameshifted protein is expressed and evades NMD.

Lastly, we report whether the truncation or resulting frameshifted protein affects known Pfam protein domains. This will occur if the domain intersects with the exitron sequence or is downstream of an exitron induced frameshift.

### S3 Benchmarking Procedures

To simulate exitrons, we chose 10,000 protein coding transcripts from the GENCODE v37 annotation at random with replacement. For each transcript we chose a random CDS region and found random G[T/C]-AG splice sites at least 30 nt away from each other (with a 9:1 GT to GC splice-site ratio). We then saved the resulting spliced RNA sequence in FASTA format. We then used PBSIM2[2] to simulate reads from each transcript at a depth of 100 with the Oxford Nanopore (ONT) R95 chemistry HMM. Because we are interested in direct RNA ONT sequencing, we simulated only positive-sense reads rather than both positive and negative-sense reads, as one would get using, for example, cDNA library preparation. We thus made some slight adjustments to the PBSIM2 simulator, which can be found at: https://github.com/ylab-hi/pbsim2. The simulated reads were aligned to the reference genome using Minimap2 v2.17.[3]

We repeated this procedure setting PBSIM2's mean read accuracy at 80%, 85%, 90%, 95% and 100%, resulting in 50,000 simulated exitrons. We ran ScanExitronLR on these reads

---

[1] Lindeboom, Rik G H et al. "The rules and impact of nonsense-mediated mRNA decay in human cancers." *Nature genetics* vol. 48,10 (2016): 1112-8. doi:10.1038/ng.3664

[2] Ono, Yukiteru et al. "PBSIM2: a simulator for long-read sequencers with a novel generative model of quality scores." *Bioinformatics (Oxford, England)* vol. 37,5 (2021): 589-595

[3] Li, Heng. "Minimap2: pairwise alignment for nucleotide sequences." *Bioinformatics (Oxford, England)* vol. 34,18 (2018): 3094-3100.

and compared the output to the ground-truth. To evaluate the performance of ScanExitronLR, we also ran ScanExitron, a published tool designed to detect exitrons in short-read RNA-seq data.[4] However, the current implementation of ScanExitron is incompatible with Minimap2 alignments--short-read aligners like HISAT2 and STAR report the read's originating strand using the XS supplementary tag, which ScanExitron depends on. However, Minimap2 reports the read's originating strand using a related but not directly equivalent ts:A tag. A small processing step of Minimap2's BAM alignment output can add the requisite XS tag. After this processing step, we ran ScanExitron on the simulated reads and compared them to the groud-truth.

In addition to simulating exitron spliced reads, we also simulated unspliced reads from every gene transcript at a depth of 100 with 100% read accuracy and ran ScanExitronLR on the dataset to see if our algorithm would produce false positives. Surprisingly, we found 43 genes that contained exitrons that were supported by at least two reads. All these false spliced alignments arose from highly repetitive regions. We include these regions in a blacklist and they are automatically filtered out in our published tool.

In order to benchmark our algorithm on real long-read datasets, we compared long-read exitron quantification predicted by ScanExitronLR with short-read predictions by ScanExitron on direct RNA ONT data from three cancer cell lines from the Singapore Nanopore Expression Project (SGNex v0.2) and ScanExitron on Illumina short-read replicates (Chen et al, 2021). We downloaded five replicate raw direct RNA FASTQ files from HepG2 (ID: ERR6053071, ERR6053072, ERR6053073, ERR6053074, ERR6053076), three replicate raw direct RNA FASTQ files from A549 (ID: ERR6053024, ERR6053025, ERR6053027) and four replicate raw direct RNA FASTQ files from MCF7 (ID: ERR6053099, ERR6053100, ERR6053101, ERR6053103). For each cell line, we merged the replicate alignments and aligned them to the human reference genome GRCh38 using Minimap2 v2.17 with recommended direct RNA settings. For short-read Illumina data, we downloaded three replicate raw FASTQ files from HegG2 (ID: ERR6937791, ERR6937792, ERR6937793), three replicate raw FASTQ files from A549 (ID: ERR6937788, ERR6937789, ERR6937790) and three replicate raw FASTQ files from MCF7 (ID: ERR6937800, ERR6937801, ERR6937802). For each cell line, we aligned each FASTQ file to the human reference genome GRCh28 using STAR 2.7.10a with default parameters. For each alignment we ran ScanExitronLR and ScanExitron with the GENCODE v37 annotation reference and a PSO cuttoff of 0.05 and AO cuttoff of 2. In addition, because low coverage genes in direct RNA may lead to particularly noisy PSO measurements, we only compared exitrons with AO/PSO > 20.

We tested how long-read exitron PSO compares with short-read exitron PSO by running ScanExitronLR on the direct RNA ONT data and ScanExitron on Illumina short-read replicates. We compared each direct RNA sample to each short-read replicate, looking for exitrons found in both pairs. Of the exitrons found in both short-read and long-read data, PSO values are strongly correlated (Pearson's correlation coefficient $r = 0.758$, $p = 1.9\text{x}10^{-10}$; Supplementary Figure S3b). We also ran ScanExitron on the direct RNA ONT data and similarly looked for exitrons found in both direct RNA and short-read pairs. We found that PSOs were less correlated and, moreover, were skewed toward higher PSOs in the short-read

[4] Wang, Ting-You et al. "A pan-cancer transcriptome analysis of exitron splicing identifies novel cancer driver genes and neoepitopes." *Molecular cell* vol. 81,10 (2021): 2246-2260.e12. doi:10.1016/j.molcel.2021.03.028

Illumina data (Pearson's correlation coefficient $r = 0.687$, $p = 3.6 \times 10^{-7}$; Supplementary Figure S2c). This is consistent with our simulation data, which showed that the short-read algorithm missed reads harboring exitrons in the long-read data and reported lower PSOs compared to the ground-truth.

**S4 Arabidopsis Example**

*Aradopsis Thaliana* raw direct RNA fastq files were downloaded from GEO accession GSM4298910 and aligned to the TAIR10 reference genome using Minimap2 v2.17 using recommended direct RNA settings. Exitrons were detected using ScanExitronLR with the TAIR10 reference gene annotation and an AO cutoff of 2 and PSO cutoff of 0.05. A detailed explanation of how to recreate this example is located at our github page (https://github.com/ylab-hi/ScanExitronLR/tree/main/test_data).

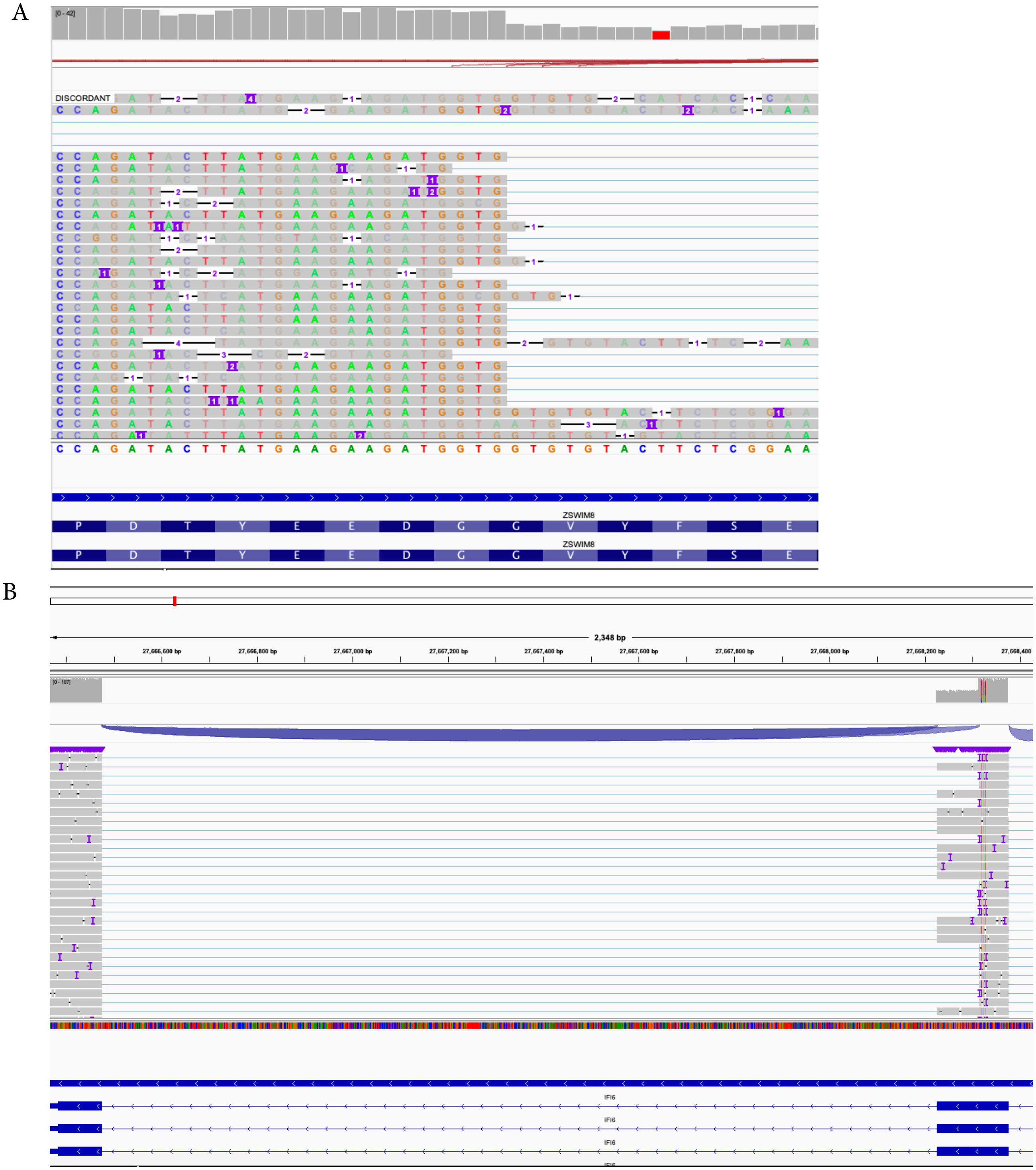


**Supplementary Figure 1.** (A) IGV view of exitron splicing event in HepG2 direct RNA sequencing sample. Some of the exitron splice sites are misaligned due to the presence of nearby GT sequences and insertions and deletions. (B) IGV view of alternative 3' splice-site misalignments in simulated data.

A

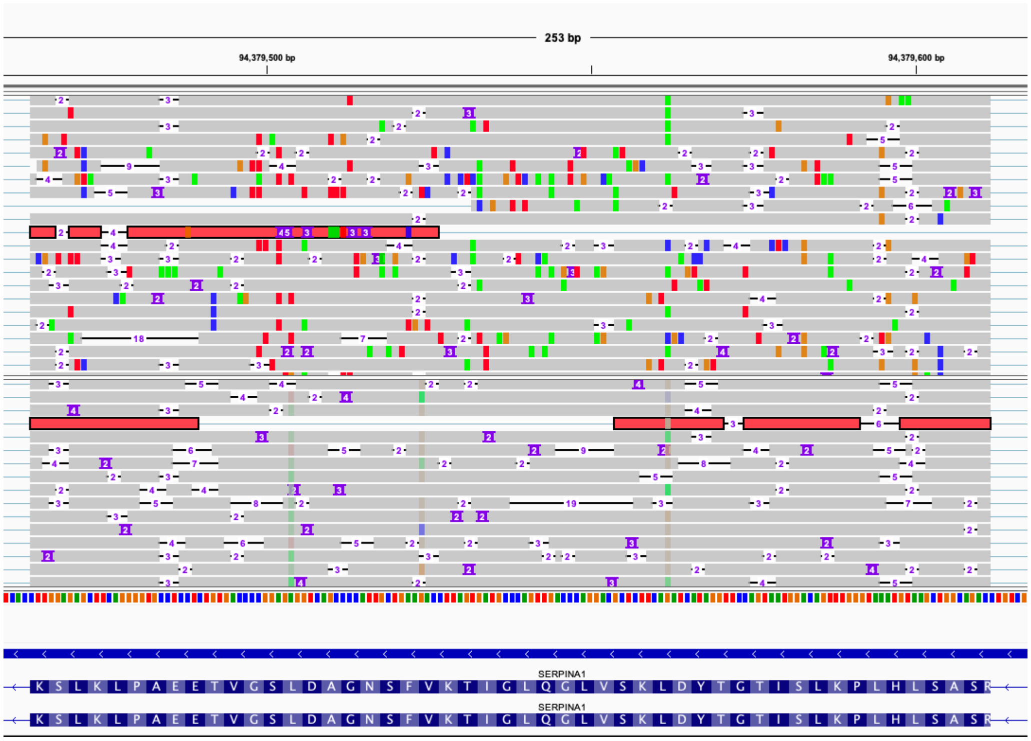


B

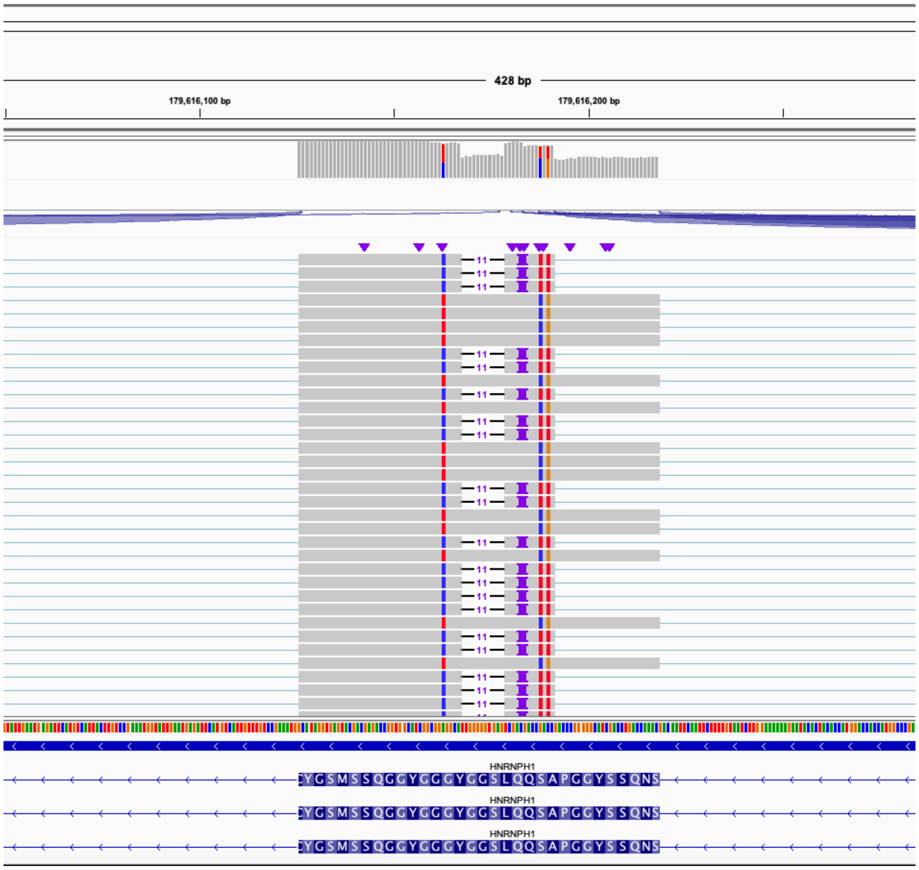


**Supplementary Figure 2**. (A) IGV view of exitron splicing event in HepG2 direct RNA sequencing sample. The second read highlighted in red harbors the exitron, while the first read highlighted in red has been realigned to the exitron splicing event. Note the many insertions (of length 4, 5, 3, 3, and 3) clumped within the misaligned exitron region. (B) IGV view of simulated exitron not detected by ScanExitronLR at 100% mean read accuracy. Thus, even at 100% read accuracy misaligned exitrons can occur.

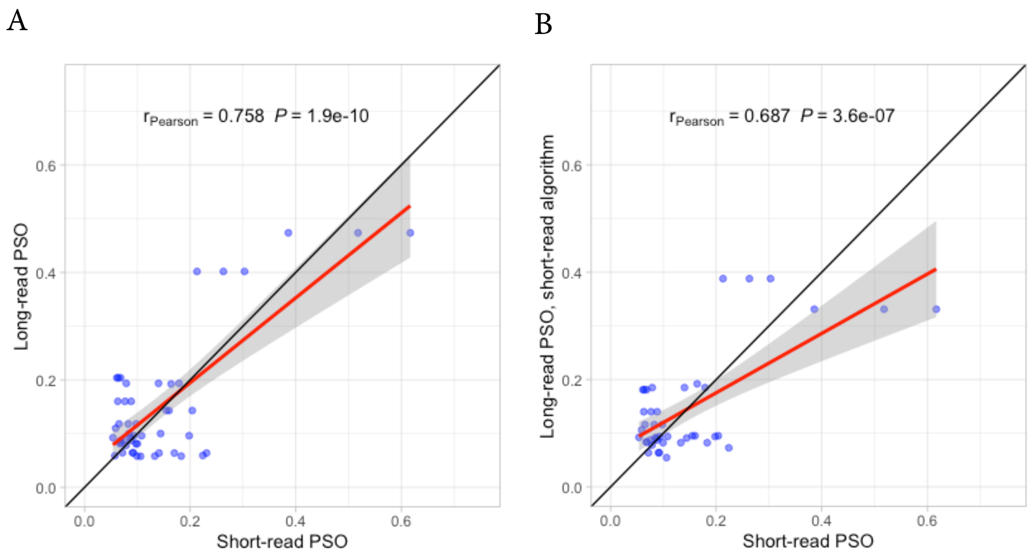


**Supplementary Figure 3**. (A) Correlation between long-read exitrons found in direct RNA ONT sequencing and short-read Illumina sequencing of MCF7, A549 and HepG2 cell lines from SGNex v0.2. (B) Same correlation but using short-read algorithm to call exitrons in the direct RNA ONT data.
